# The geometry of normal tissue and cancer gene expression manifolds

**DOI:** 10.1101/2021.08.20.457160

**Authors:** Joan Nieves, Augusto Gonzalez

## Abstract

A recent paper shows that in gene expression space the manifold spanned by normal tissues and the manifold spanned by the corresponding tumors are disjoint. The statement is based on a two-dimensional projection of gene expression data. In the present paper, we show that, for the multi-dimensional vectors defining the centers of cloud samples: 1. The closest tumor to a given normal tissue is the tumor developed in that tissue, 2. Two normal tissues define quasi-orthogonal directions, 3. A tumor may have a projection onto its corresponding normal tissue, but it is quasi-orthogonal to all other normal tissues, and 4. The cancer manifold is roughly obtained by translating the normal tissue manifold along an orthogonal direction defined by a global cancer progression axis. These geometrical properties add a new characterization of normal tissues and tumors and may have biological significance. Indeed, normal tissues at the vertices of a high-dimensional simplex could indicate genotype optimization for given tissue functions, and a way of avoiding errors in embryonary development. On the other hand, the cancer progression axis could define relevant pan-cancer genes and seems to be consistent with the atavistic theory of tumors.

## 1 Introduction

In gene expression (GE) space, the normal homeostatic and cancer phenotypes for a given tissue span distant regions allowing an easy classification of samples (Alon et al., 1999; Gonzalez et al., 2023). A similar situation takes place in protein expression (PE) space (Jane et al., 2001). The transition between these two radically different states of the tissue is understood to proceed discontinuously, through the rearrangement of the expressions of thousands of genes (Gonzalez et al., 2021).

On the other hand, different normal tissues in the human body are also well separated in GE or PE spaces. We understand that this separation has an epigenetic origin (Zhang et al., 2013). The set of all normal tissues span a normal tissue manifold, which experiences only minor changes during the individual lifespan because of its homeostatic nature.

In Gonzalez et al. (2023), it was shown that the set of cancer attractors also span a manifold in GE space, well apart from the normal tissue manifold. The atavistic origin hypothesis for tumors (Davies and Lineweaver, 2011) was used to argue that they should be grouped. A main direction indicating cancer progression was defined. This direction is probably related to genes playing an important role in any cancer. The statement is based on a two-dimensional representation of gene expression data. In the present paper, we elaborate on the multi-dimensional geometry of normal tissue and cancer manifolds in GE space. Previously, we have shown that geometric magnitudes are biologically meaningful. For example, the distance between two tumor centers shows an inverse dependence on the number of common differentially expressed genes (Gonzalez et al., 2023). Here, we look for a more detailed geometrical representation of both manifolds, that is, to answer questions like which are the actual tumors closest to a given normal tissue, which is the true direction of the cancer projection axis, etc.

We process RNA-seq GE data for 15 tissues and their corresponding tumors, obtained from The Cancer Genome Atlas portal (TCGA) (Tomczak et al., 2015). For the vectors defining the centers of the clouds of normal and tumor samples in each tissue, the following statements are proven: 1. The closest tumor to a given normal tissue is the tumor developed in that tissue, 2. Two normal tissues define quasi-orthogonal directions, 3. A tumor may have a projection onto its corresponding normal tissue, but it is quasi-orthogonal to all other normal tissues, and 4. The cancer manifold is roughly obtained by translating the normal tissue manifold along an orthogonal direction defined by a global cancer progression axis.

In summary, normal tissues seem to be located at the vertices of a nearly regular simplex or polytope. For single cell gene expression data, the vertices of 2-, 3– and 4-polytopes are related to genotypes optimizing specific cellular functions (Korem et al., 2015). This interpretation is probably valid for our tissue data. Below, we shall further speculate on the biological implications of the geometric properties of manifolds.

The plan of the paper is as follows. After a brief declaration of variables and methods, the main results of the paper are listed in Section 2. Section 3 contains the conclusions. Dimensional reduction results for protein expression data are given in the Appendix in order to show that normal tissues and tumor manifolds are also separated in protein expression space.

## 2 The normal tissue and cancer manifolds

First, let us define the variables. In Table 1, we list the studied tissues and the number of normal and tumor samples in each one. The data includes the expressions of 60483 genes, thus each sample is defined by an expression vector of dimension 60483. This is the typical situation in gene– and protein-expression data analysis (Clarke et al., 2008), where the number of genes is much larger than the number of samples. The *i*-component of the sample vector is given by:

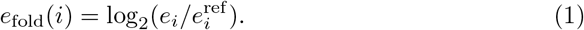

**Table 1.** The TCGA abbreviations and the number of samples in the 15 tissues under study.

| Cancer type | Abbreviation | Normal samples | Tumor samples |
| --- | --- | --- | --- |
| Bladder urothelial carcinoma | BLCA | 19 | 414 |
| Breast invasive carcinoma | BRCA | 112 | 1096 |
| Colon adenocarcinoma | COAD | 41 | 473 |
| Esophageal carcinoma | ESCA | 11 | 160 |
| Head and neck squamous cell carcinoma | HNSC | 44 | 502 |
| Kidney clear cell carcinoma | KIRC | 72 | 539 |
| Kidney papillary cell carcinoma | KIRP | 32 | 289 |
| Liver hepatocellular carcinoma | LIHC | 50 | 374 |
| Lung adenocarcinoma | LUAD | 59 | 535 |
| Lung squamous cell carcinoma | LUSC | 49 | 502 |
| Prostate adenocarcinoma | PRAD | 52 | 499 |
| Rectum adenocarcinoma | READ | 10 | 167 |
| Stomach adenocarcinoma | STAD | 32 | 375 |
| Thyroid carcinoma | THCA | 58 | 510 |
| Uterine corpus endometrial carcinoma | UCEC | 23 | 552 |

The variable *e*_fold_(*i*) gives the logarithmic fold variation of gene *i* with respect to the reference level 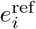. We shall use the same reference for all tissues, constructed by geometric averaging over all normal samples. The method is described elsewhere (Gonzalez et al., 2023). A value *e*_fold_(*i*) *>* 0 means that the gene is over-expressed with regard to the reference, whereas *e*_fold_(*i*) *<* 0 means that the gene is under-expressed or silenced.

From sample vectors, we define tissue vectors by averaging over tissue normal or tumor samples, respectively: **e**_tissue_(normal), and **e**_tissue_(tumor). They may be understood as the centers of sample clouds. Indeed, a Principal Component Analysis (PCA) (Lever et al., 2017) of the data for liver hepatocellular carcinoma (LIHC), for example, shows a 2D schematic representation of samples in GE space (Gonzalez et al., 2023), as it is apparent in Fig. 1. Cloud centers are well defined. We shall work with these centers as tissue representatives.

**Fig. 1.**
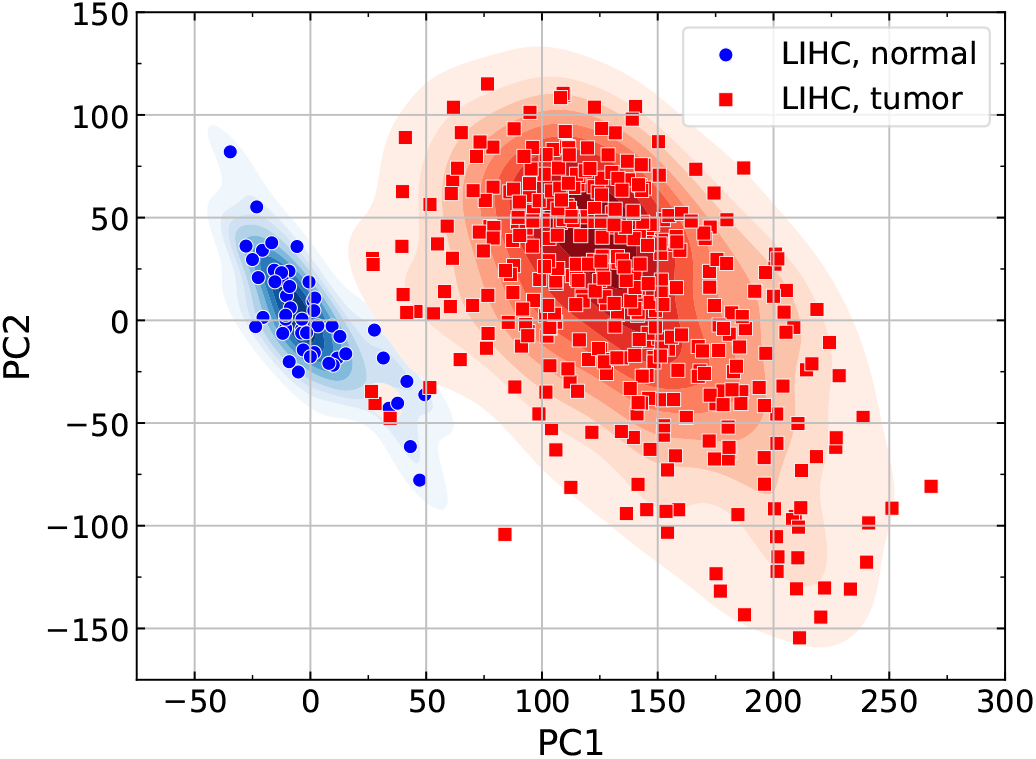
Reduced dimensionality representation of normal and tumor samples in LIHC. A density plot of samples is added with the purpose of easy visualization of the cloud centers.

In Gonzalez et al. (2023), we show in a 2D scheme that tumor centers define a tumor manifold, well separated from the normal tissue manifold. A 3D representation, obtained from the PCA of all studied tissues, is given in Fig. 2. A schematics of results is drawn in the right panel. Manifolds are well separated and a cancer progression axis may be defined. We shall show that this global picture is correct, although actual distances or angles are not properly represented in this reduced dimensionality diagram.

**Fig. 2.**
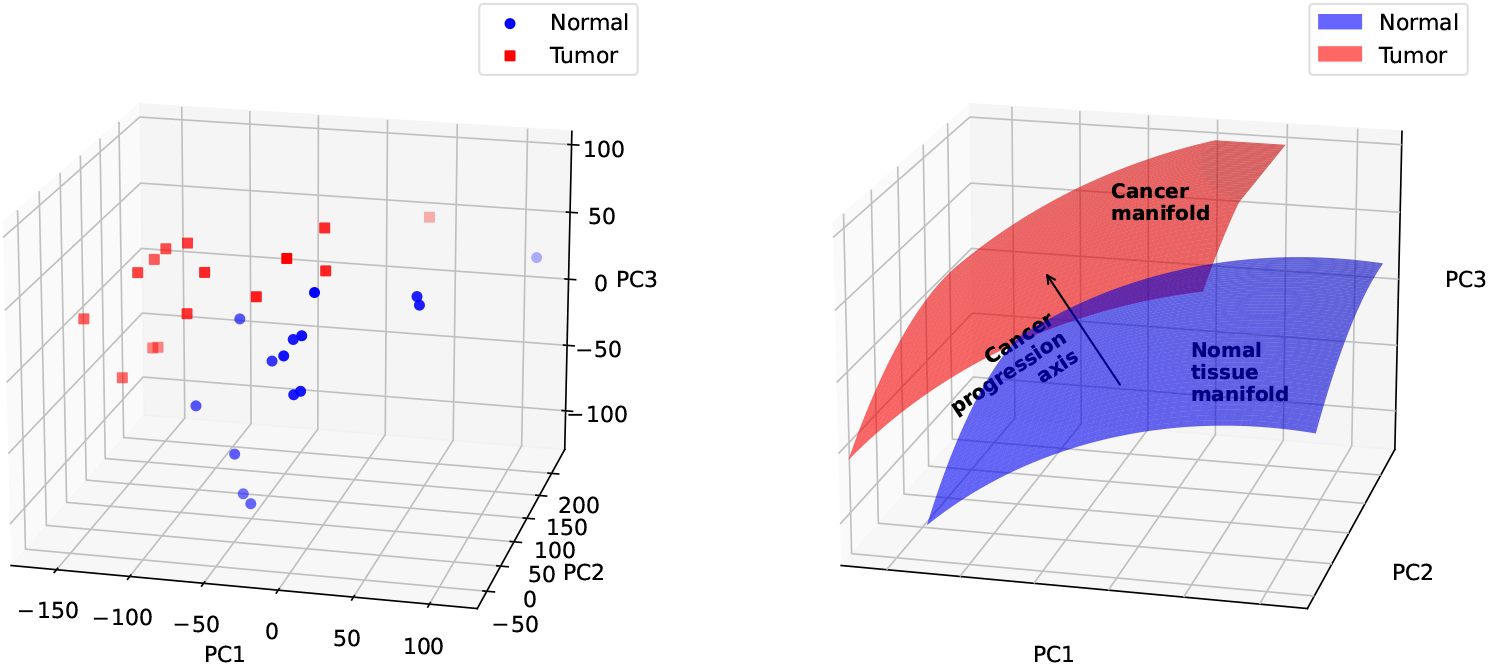
Left: The first three principal components for TCGA gene expression data for 15 tissues. The centers of the clouds of samples for each tissue are depicted. Right: Schematics of the topology of gene expression space. The sets of normal tissues and cancer attractors are ideally represented as spanning smooth manifolds, with a cancer progression axis connecting the centers of both regions.

A simple way to define the cancer progression axis could be the vector that connects the centers of both manifolds. We define it as follows:

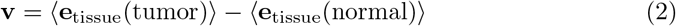

Where ⟨… ⟩means average over tissues.

We first show that all vectors connecting a normal tissue to its respective tumor have a positive projection onto the cancer progression axis. In other words, the schematics shown in Fig. 2 is preserved in higher dimensions. The exact formulation of this statement is the following:

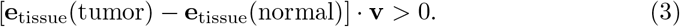

The results for the normalized projections, i.e. divided by the norms of both vectors, are listed in the second column of Table 2.

**Table 2.** Some geometric properties of the normal tissues and cancer manifolds. 2nd column: The normalized projections over the cancer progression axis of vectors connecting a given normal tissue to its tumor, Eq. (3). 3rd column: The angle between a normal tissue vector and the cancer progression axis, *θ*(N)*/π*. 4th column: The distance between a given normal tissue and its tumor. The closest tumor, if different from the proper one, is indicated in parenthesis.

| Tissue | Normalized projection | $\theta(N)/\pi$ | dist(N,T) |
| --- | --- | --- | --- |
| BLCA | 0.80 | 0.48 | 140.97 |
| BRCA | 0.65 | 0.51 | 137.53 |
| COAD | 0.68 | 0.48 | 156.05 |
| ESCA | 0.67 | 0.43 | 139.69 (136.77, STAD) |
| HNSC | 0.61 | 0.43 | 123.57 |
| KIRC | 0.42 | 0.56 | 173.02 (169.91, KIRP) |
| KIRP | 0.52 | 0.54 | 163.58 |
| LIHC | 0.51 | 0.52 | 135.72 |
| LUAD | 0.68 | 0.54 | 147.34 |
| LUSC | 0.67 | 0.51 | 194.99 (142.52, LUAD) |
| PRAD | 0.50 | 0.49 | 92.68 |
| READ | 0.71 | 0.50 | 168.13 (166.22, COAD) |
| STAD | 0.71 | 0.48 | 137.46 |
| THCA | 0.32 | 0.49 | 113.00 |
| UCEC | 0.67 | 0.51 | 171.43 |

### 2.1 The closest tumor to a given normal tissue is its tumor

As mentioned above, actual distances are not properly represented in 2D or 3D PCA projections. The distance from a normal tissue N to a tumor T^*′*^ is defined as:

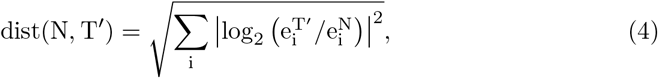

where the sum runs over genes, and 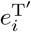 and 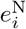 are, respectively, tissue expressions of gene *i* in T^*′*^ and N. Thus, the distance is an *l*^2^-norm of the log-fold differential expressions when N is taken as reference.

The distances between a given normal tissue and its tumor, computed from Eq. (4), are given in the fourth column of Table 2. It shows that, in general, the closest tumor to a given normal tissue is its tumor. Exceptions to this rule are tumors in the same organ (KIRC – KIRP, LUSC – LUAD, READ – COAD), and the ESCA – STAD pair, which is not in the same organ.

This statement may be rephrased as follows: Differential expressions for a tumor are the smallest when the corresponding normal tissue is taken as reference.

Notice that distances are normally distributed around the mean, 146.34, that is 73 % of the distances (11 from 15) are closer than one standard deviation from the mean.

### 2.2 A pair of normal tissues define quasi-orthogonal directions in GE space

We shall compute the angles *θ*_NN_*′* defined from the scalar product between two normal tissue vectors, i.e.

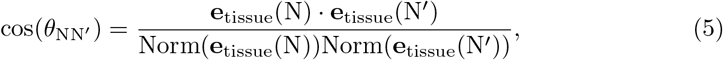

where Norm is the euclidean norm of the vector. Figure 3a) shows an histogram of *θ*_NN_*′* values. Angles are distributed around 0.50 *π* with standard deviation 0.09 *π*. Deviations 5 are related to the three pairs of tissues in the same organ, and the pair esophagus – stomach, mentioned above.

**Fig. 3.**
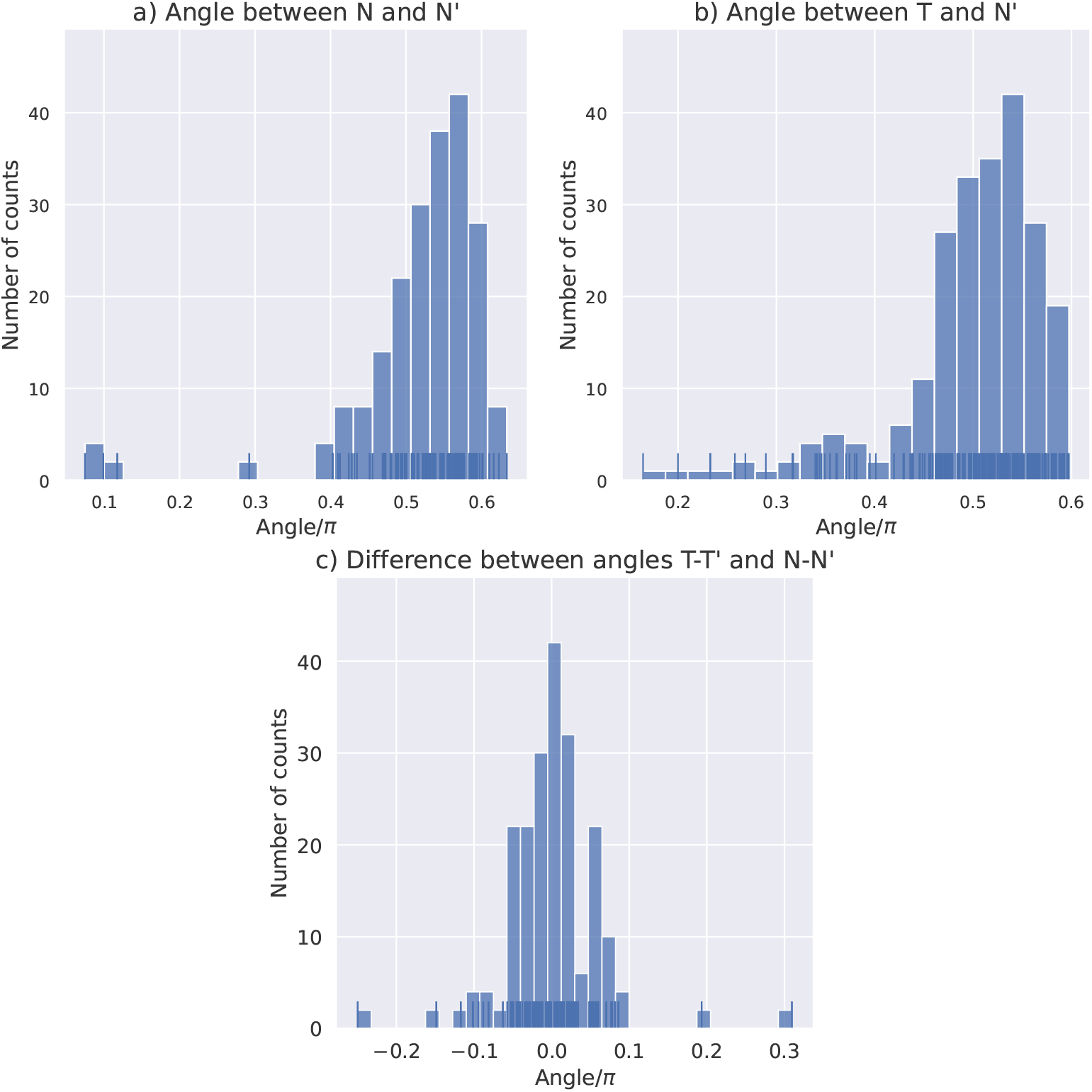
a) Histogram for the angle between two normal tissue vectors. The average value for the angle in both panels a) and b) is 0.50 *π*. b) Histogram for the angle between tissue vectors corresponding to T and N^*′*^. c) Histogram for the difference between the angles *θ*_TT_*′* and *θ*_NN_*′*, measured in their respective manifolds. The average value for this difference is 0.001 *π*.

Thus, two normal tissue vectors define quasi-orthogonal directions. The meaning of such a statement is as follows: gene expressions in two normal tissues are not correlated. That is, genes expressed in the first tissue are not related to those expressed in the second one.

It is interesting to compute also the norm of normal tissue vectors, that is the distance from the global reference to the center of the normal sample cloud in a given tissue. All normal tissues radii are within one standard deviation from the mean: 154.63 ± 36.06. The exceptions are BLCA and LIHC, for which Norm(BLCA) = 109.29, and Norm(LIHC) = 270.80. These are the closest and the farthest from the global reference, respectively.

According to these results we may say that the normal tissue vectors span a manifold similar to a regular simplex or polytope, where pairs of vectors are quasi orthogonal. We shall come back below to the interpretation of this geometrical property.

### 2.3 A tumor tissue vector may have non-zero projection onto its normal tissue, but it is quasi-orthogonal to the rest

The mean value of angles *θ*_TN_*′* between tissue vectors corresponding to the tumor T and the normal tissue N^*′*^ is again 0.50 *π*, with standard deviation 0.08 *π*. The results are drawn in Fig. 3b).

In general, we may say that there are non-zero projections of the expression vector associated to the tumor T onto its corresponding normal tissue N and other tissues in the same organ, but this vector is quasi-orthogonal to the rest of normal tissues.

One may check also whether the angle *θ*_TT_*′* between a pair of tumor tissue vectors, measured in the cancer manifold, is similar to the angle *θ*_NN_*′* between the pair of corresponding normal tissues. In Fig. 3c) we draw the histogram for the difference. Indeed, angles are roughly preserved between tumor and normal tissue pairs of vectors. For the mean difference, we obtain: (0.001 ± 0.056) *π*. The outlayers are related to tumors in the same organ.

A final question concerns the orthogonality of the cancer progression axis, **v**, to the normal tissue manifold. As mentioned above, in the third column of Table 2, we show the projections onto each of the normal tissue vectors. It is apparent that the cancer progression axis roughly defines an orthogonal direction. For the mean angle, we obtain 0.50 *π*.

In general, we may say that tumor vectors are also distributed on a sector of a hyper-surface, which is roughly obtained by translating the normal tissue manifold along the orthogonal direction defined by the cancer progression axis.

## 3 Conclusion

We use the vectors defining cloud centers of GE data in order to compute geometrical properties of normal tissue and tumor manifolds. The 3D schematic representation given in Fig. 2, right panel, is shown to hold in higher dimensions. Roughly speaking, normal tissues are evenly distributed on a hyper-spherical surface. Any pair of vectors is quasi-orthogonal. In other words, they are located at the vertices of a regular polytope. Tumors also lie on a manifold, which is roughly obtained by translating the normal tissue manifold along the cancer progression axis.

We noticed that the geometry of normal tissue manifold shares similarities with the geometry of the manifold for single cells realizing multiple tasks (Korem et al., 2015). In that case, the vertices of the polytope are related to genotypes optimizing a single cellular function. It is thus natural that tissues, which usually realize very specific functions, localize themselves at the vertices of their polytope. On the other hand, at a more speculative level, the quasi-orthogonality of normal tissues could be related to preserving the identity or avoiding errors in embryonic development. In other words, most genes are tissue specific and the probability of rising by chance genes not related to the tissue in embryonic development is very low. An additional supporting evidence in this respect is the quasi-orthogonality of the set of genes realizing circadian oscillations in different tissues, the cyclers (Ruben et al., 2018).

It is interesting to speculate further and look at these geometrical results from the point of view of the atavistic theory of tumors. Cooperation between unicellular organisms leading to primitive metazoans could probably be related to a process of mixing and orthogonalization of their genome as a way of optimizing specific functions. This hypothesis could perhaps be checked in the simplest multicellular organisms. The evolution towards advanced pluricellularity could then proceed through an orthogonal direction in GE space, in a direction opposite to the cancer progression axis.

The geometry of GE data provides an additional characterization of normal tissues and tumors, for which there are other measurements already available. For example, the degree of disorder may be described by the entropy of the expression profile (Mesa-Rodríguez et al., 2022). Tumors are shown to be more entropic than normal tissues. A related measurement comes from the estimation of the volume of the basins of attraction for both normal and tumor attractors in GE space (Gonzalez et al., 2022). The number of tumor genotypes is shown to be exponentially greater than that of normal tissue genotypes. Work along these directions is in progress.

### Appendix A The normal tissue and cancer manifolds in PE space

For the sake of consistency, we show separation of the normal tissue and cancer manifolds in protein expression space.

Protein expression data coming from immuno-histo-chemistry measurements are available in the Human Protein Atlas (HPA) database (Uhlen et al., 2017). We use the normal tissue and pathology data.

Expression values are classified in one of four categories: High, Medium, Low and Not Detected. We assume that these categories may be translated into three values of the *e*_*fold*_ variable: +1, 0, –1, and –1, respectively. When various lectures are provided for a single tissue, we take the average.

A total of 10166 proteins are consistently expressed in 49 normal tissues and 20 tumors. This is the dimension of the covariance matrix in the PCA. The first three principal components emerging after diagonalization account for 41 % of the total variance. They are used to draw Figure A1.

**Fig. A1.**
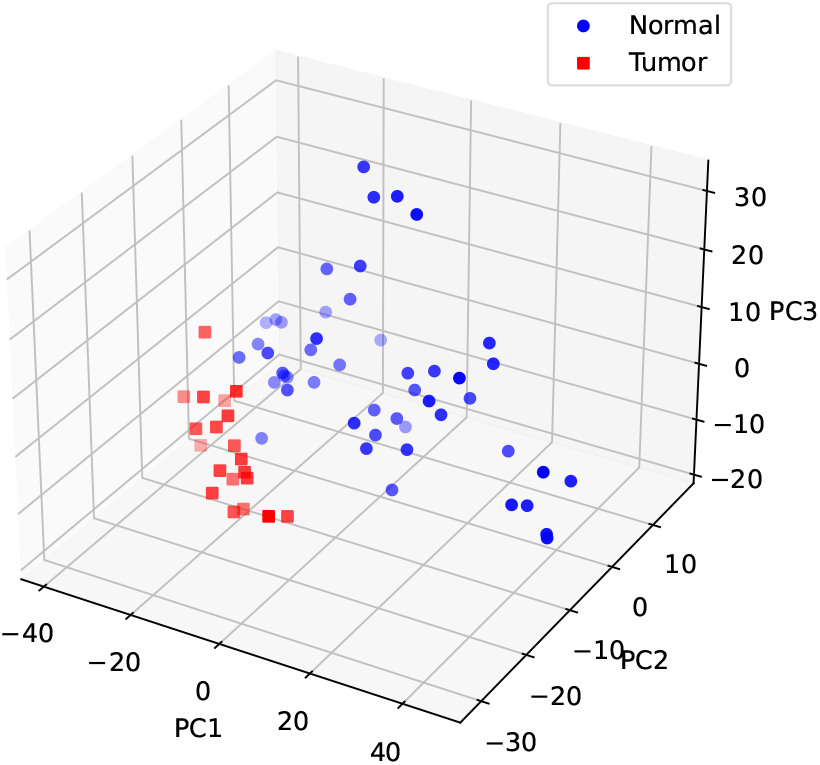
PCA results for the protein expression data of the HPA (49 normal tissues, 20 tumors). As in Fig. 2, normal tissue and cancer regions are apparent.

The tissue differentiation and cancer manifolds are also apparent in PE space. They are mainly separated along the PC2 direction. The difference with regard to GE data could be related to the discrete nature of the PE data and also to regulation of translation.

## Acknowledgments

The authors acknowledge the Cuban Agency for Nuclear Energy and Advanced Technologies (AENTA) and the Office of External Activities of the Abdus Salam Centre for Theoretical Physics (ICTP) for support.

## Authors contributions

A.G. conceived and coordinated the work. J.N. processed GE and PE data and actualized the GitHub repository. Both authors analyzed and interpreted the results, contributed to the manuscript and approved the final version.

## Availability of data and materials

The information about the data we used, the procedures and results are integrated in the public repository: https://github.com/JoanANievesCuadrado/manifolds-geometry/.

The TCGA data for two of the studied tissues and the HPA data is replicated in …/databases external. We include in the repository’s root Python scripts in order to pre-process this data. The generated mean values for each tissue are given in …/databases generated.

Scripts which perform Principal Component Analysis and generate figures and tables are also provided. These figures and tables, which are the main results of the paper, are contained in …/Figures and Tables.

## Declarations

### Conflict of interest

The authors have no financial interest or other types of conflict of interest related to the work of this article.

### Ethical approval

The authors guarantee that no study-specific approval from an ethics committee is needed for this work.

### Informed consent

The authors guarantee that no informed consent is needed for this work.

## Notes

https://github.com/JoanANievesCuadrado/manifolds-geometry

### Competing Interest Statement

The authors have declared no competing interest.

### Summary of Updates

Changed article formatting, redone all images for better visualization, added reference supporting quasi-orthogonality of normal tissue vectors, moved dataset to another repository, fixed some grammatical and spelling mistakes.

